# Harnessing methylotrophs as a bacterial platform to reduce adverse effects of the use of the heavy lanthanide gadolinium in magnetic resonance imaging

**DOI:** 10.1101/2021.06.12.448192

**Authors:** Nathan M. Good, Harvey Lee, Emily R. Hawker, Assaf A. Gilad, N. Cecilia Martinez-Gomez

## Abstract

Gadolinium is a key component of magnetic resonance imaging contrast agents that are critical tools for enhanced detection and diagnosis of tissue and vascular abnormalities. Untargeted post-injection deposition of gadolinium *in vivo*, and association with diseases like nephrogenic systemic fibrosis, has alerted regulatory agencies to re-evaluate their widespread use and generated calls for safer gadolinium-based contrast agents (GBCAs). Increasing anthropogenic gadolinium in surface water has also raised concerns of potential bioaccumulation in plants and animals. Methylotrophic bacteria can acquire, transport, store and use light lanthanides as part of a cofactor complex with pyrroloquinoline quinone (PQQ), an essential component of XoxF-type methanol dehydrogenases (MDHs), a critical enzyme for methylotrophic growth with methanol. We report robust gadolinium-dependent methanol growth of a genetic variant of *Methylorubrum extorquens* AM1, named *evo*-HLn, for “evolved for heavy lanthanides”. Genetic adaptation of *evo*-HLn resulted in increased *xox1* promoter and XoxF MDH activities, transport and storage of Gd^3+^, and augmented biosynthesis of PQQ. Gadolinium-grown cells exhibited a shorter T1 relaxation time compared to cells with lanthanum or no lanthanide when analyzed by MRI. In addition, *evo*-HLn was able to grow on methanol using the GBCA Gd-DTPA as the sole gadolinium source, showing the potential of this strain for the development of novel GBCAs and gadolinium recovery from medical waste and/or wastewater.

---

Gadolinium (Gd^3+^; atomic number 64) is a versatile element that is widely used in various modern industries (Ebrahimi and Barbieri 2019) but is perhaps best-known for its use as a contrast agent for MRI. Its seven unpaired electrons give Gd^3+^ unparalleled paramagnetic properties, making it the most effective agent for clinical application (Srivastava et al. 2015). Gd^3+^ alone is highly toxic to humans (Le Fur and Caravan 2019) and is therefore injected as a nine-coordinate ion chelated by an octadentate polyaminocarboxylate ligand with a water coligand (Wahsner et al. 2019) known as GBCAs. The stability of GBCAs makes them highly effective for intravenous delivery, and as a result they are used in an estimated 30 million MRI exams annually (Lohrke et al. 2016), with around half a billion doses administered thus far (McDonald et al. 2018). GBCAs are excreted in urine post-injection, however, they are not innocuous. Over the past two decades, the development of nephrogenic systemic fibrosis (NSF) has been observed in GBCA injection patients with impaired renal function, resulting in joint pain, immobility and even death (Grobner 2006; Marckmann et al. 2006; Boyd, Zic, and Abraham 2007; High et al. 2007). The last five years have generated rising alarm over the use of GBCAs with long-term retention found in patients with normal kidney function (Kanda et al. 2014, 2015; McDonald et al. 2015; Roberts et al. 2016). Anaphylactic shock and kidney failure have also been reported as possible outcomes of Gd^3+^ accumulation in tissues (Ergün et al. 2006; Hasdenteufel et al. 2008). Unmetabolized, excreted GBCAs are also cause for concern as rising anthropogenic Gd^3+^ in surface water correlates with steadily increasing annual MRI exams worldwide (Ebrahimi and Barbieri 2019). Due to the toxicity and rising concentrations of this microcontaminant, the potential health impacts on aquatic life and bioaccumulation in the food-chain deserve more attention, as do wastewater treatment strategies that are sufficient to remove Gd^3+^.

Gadolinium is a member of the lanthanide series of elements, a group that has recently been added as life metals. A broader understanding of the functions of lanthanides (Ln^3+^) in biology is slowly unfurling with discoveries of novel enzymes, metabolic pathways, and organisms that are dependent on these metals. Ln^3+^ are known to form a cofactor complex with the prosthetic group PQQ for some alcohol dehydrogenase enzymes (Keltjens et al. 2014). XoxF MDH from the methylotrophic bacterium *Methylorubrum* (formerly *Methylobacterium*) *extorquens* AM1 was the first reported Ln^3+^-dependent metallo-enzyme, and members of this diverse enzyme class are wide-spread in marine, fresh water, phyllosphere, and soil habitats (Nakagawa et al. 2012; Taubert et al. 2015; Huang, Yu, and Chistoserdova 2018; Chistoserdova 2016; Ochsner et al. 2019; Keltjens et al. 2014). ExaF ethanol dehydrogenase was the first reported Ln^3+^-dependent enzyme with a preference for a multi-carbon substrate, and its discovery has led to the identification of related enzymes in non-methylotrophic bacteria (Wehrmann et al. 2017; Wegner et al. 2019). Ln^3+^ are also known to influence metabolic pathways in methylotrophic and non-methylotrophic bacteria (Good et al. 2019; Wehrmann et al. 2020). To date, all known Ln^3+^-dependent metallo-enzymes are from bacteria and coordinate the metal-PQQ complex for catalytic function. However, the physiological importance of PQQ stretches well-beyond the prokaryotes. Mammals, including humans (Killgore et al. 1989), and plants (Choi et al. 2008) benefit from PQQ. Eukaryotes (Takeda et al. 2015; Matsumura et al. 2014) and archaea (Sakuraba et al. 2010) produce PQQ-dependent enzymes, though there is still much to be discovered regarding their activities and function. Nonetheless, the link between PQQ and Ln^3+^-dependent metallo-enzymes suggests that the influence of Ln^3+^ in biological processes may spread across all three Kingdoms of life.

Evidence for the biological use of Ln^3+^ in bacteria was first reported as the stimulation of methanol growth and expression of PQQ-MDH activity in bacterial cultures grown with lanthanum (La^3+^; atomic number 57) or cerium (Ce^3+^, atomic number 58) (Hibi et al. 2011; Fitriyanto et al. 2011). At the time, Ln^3+^ were considered unavailable and unutilized for biological processes due to their insolubility in nature, and it was proposed that though Ln^3+^ are more potent Lewis acids than calcium (Ca^2+^), evolution likely passed them by in favor of the more bioavailable metal (Lim and Franklin 2004). PQQ-MDHs were typified by MxaFI, an α2β2 tetrameric enzyme that coordinates Ca^2+^ in the large subunit of each protomer (Richardson and Anthony 1992; Adachi et al. 1990). MDH is a critical enzyme for methylotrophic bacteria, organisms that can oxidize reduced carbon compounds with no carbon-carbon bonds, such as methane and methanol, and has been the subject of genetic, biochemical and chemical studies for decades (Christopher Anthony and Williams 2003; Zhang, Reddy, and Bruice 2007; Zheng et al. 2001; Goodwin and Anthony 1996; Williams et al. 2005; M. Ghosh et al. 1995). Shortly after the discovery of Ln^3+^ dependence for XoxF MDH activity in *M. extorquens* AM1 (Nakagawa et al. 2012), the extremophile methanotroph *Methylacidiphilum fumariolicum* SolV was shown to rely on Ln^3+^ in its volcanic mudpot environment for survival (Pol et al. 2014). Several subsequent studies noted the role of Ln^3+^ in regulating MDH expression (Farhan Ul Haque et al. 2015; Vu et al. 2016; Chu and Lidstrom 2016), describing the “lanthanide-switch” phenomenon in which the presence of light Ln^3+^ up-regulates expression of *xox* genes and concomitantly down-regulates expression of *mxa* genes. Global studies have suggested that Ln^3+^ may impact more than MDH and accessory gene expression, including alterations to downstream metabolism (Gu et al. 2016; Good et al. 2019; Masuda et al. 2018).

Growth studies with mesophilic methylotrophs and the Ln^3+^ series of metals have shown that only members of the “light” classification, ranging from La^3+^ to Nd^3+^ (atomic number 60), can support growth with XoxF MDH similar to Ca^2+^ with MxaFI MDH (Daumann 2019). In comparison, methanol growth with Sm^3+^ is much slower and growth has not been reported for Ln^3+^ of higher atomic numbers, with a couple of exceptions (Vu et al. 2016; Huang, Yu, and Chistoserdova 2018; Wang et al. 2019). *M. fumariolicum* SolV can grow with the light/heavy lanthanide Eu^3+^ well enough to produce cultures for enzyme purification (Jahn et al. 2018). This organism was also reported to show slow growth with Gd^3+^, but no studies have investigated this further (Pol et al. 2014). *M. fumariolicum* SolV grows optimally in acidic conditions (pH 2-5) making Ln^3+^ soluble for uptake and utilization, and as such does not have a known dedicated transport system for these metals. In contrast, methylotrophs that grow at neutral pH have an ABC transport system and specific TonB-dependent receptor encoded in a “lanthanide-utilization and transport” gene cluster (Roszczenko-Jasińska et al. 2020; Ochsner et al. 2019). Of such organisms known to date, only a genetically manipulated mutant strain of *Methylotenera mobilis* JLW8 has been reported to show indications of growth with Gd^3+^ in the form of increased culture density (Huang, Yu, and Chistoserdova 2018). Thus, the heavy lanthanide Gd^3+^ is the highest atomic number species known to support methanol growth in methylotrophic bacteria. Activity of XoxF MDH decreases with increasing atomic radius for the light Ln^3+^ (Jahn et al. 2018; Good et al. 2019). While decreasing XoxF MDH activity correlates with reduced growth rates seen with Ln^3+^ of increasing atomic mass, it is still not known if this is due solely to decreased enzyme catalysis or if transport of the metal ions plays a role as well. Regardless of the factor(s) limiting growth, Gd^3+^ seems to be the pivotal Ln^3+^ marking the threshold of life with these metals.

In this study we report the characterization of a *M. extorquens* AM1 genetic variant that is capable of robust growth on methanol with the heavy lanthanide Gd^3+^, a Ln^3+^ that does not support growth in the ancestral strain. We identified the variant as having a single non-synonymous substitution in a putative hybrid histidine kinase/response regulator resulting in a gain of function mutation. The gene encoding this putative regulatory system was recently identified as important to ExaF ethanol dehydrogenase Ln metabolism in *M. extorquens* AM1 (Huong N. Vu, Gabriel A. Subuyuj, Ralph Valentine Crisostomo, Elizabeth Skovran 2021). Our variant exhibited increased *xox1* promoter and MDH activities, a distinctive bright pink coloration corresponding to augmented PQQ production, and increased transport and storage of gadolinium. Accumulation of gadolinium was sufficient to generate a significant reduction in T1 relaxation time when scanned by MRI. Finally, we show that the variant could grow efficiently with the GBCA Gd-DTPA as the sole Ln^3+^ source. These discoveries provide novel avenues for bioremediation and Ln^3+^ recycling. Elucidation of the mechanisms governing Ln^3+^ uptake, storage, and usage will aid in the identification and development of genetically encoded and peptide-based imaging agents.

## RESULTS

### Isolation of an *M. extorquens* AM1 mutant strain capable of gadolinium-dependent methanol growth

The Δ*mxaF* mutant strain of *M. extorquens* AM1 can grow on methanol when provided an exogenous source of light lanthanides ranging from La^3+^ to Sm^3+^, but the heavy lanthanide Gd^3+^ had not been tested (Vu et al. 2016). We tested the ability of Δ*mxaF* to grow on methanol with Gd^3+^ as the sole Ln^3+^ available. MP methanol minimal medium with Gd^3+^ was inoculated with Δ*mxaF* and culture density was measured over time. No detectable increase in culture density was observed after 14 days of incubation at 30°C. However, after another 7 days of incubation the culture density had increased ~2.3 fold, reaching a final OD_600_ of 0.35 ± 0.03 (N = 4). Gd^3+^-grown cells were transferred to fresh methanol minimal medium with Gd^3+^ and grown to maximum culture density. This process was repeated twice.

To verify that the cultures were not contaminated, 5 μL was plated onto solid minimal succinate medium with 50 μg/mL rifamycin and incubated at 30°C (Figure S1B). Growth of pink colonies indicated the cultures were *M. extorquens* AM1, as the strain used is rifamycin-resistant (Nunn and Lidstrom 1986). Using colony PCR, we determined that cells recovered from the Gd^3+^-grown cultures were negative for *mxaF*, as was the ancestral strain, and positive for *fae* encoding formaldehyde-activating enzyme, another genetic marker specific for *M. extorquens* AM1 (Figure S1A). Cells from these Gd^3+^-grown cultures were washed four times with sterile minimal medium to remove possible residual extracellular Gd^3+^, resuspended in 1 mL sterile medium, and saved as freezer stocks with 5% DMSO at −80°C.

The long incubation time of the original cultures prior to growth with Gd^3+^ suggested either an extended period of metabolic acclimation or genomic adaptation. To discern between these two possibilities, we tested methanol growth after first passaging the strain three times on solid succinate medium and then inoculating into liquid succinate medium to generate pre-cultures. Cells from the liquid culture were harvested, washed four times with sterile minimal medium, and then inoculated into methanol medium with Gd^3+^. Growth was measured using a microplate spectrophotometer (Fig. 1A). The variant strain exhibited growth within ~15 hours of inoculation, a specific growth rate of 0.03 ± 0.00 h^-1^, and a maximum culture density 0.69 ± 0.04. The lack of the 3-week lag in growth, as we observed with the ancestral Δ*mxaF* inoculation, was indicative of genomic adaptation, rather than metabolic acclimation, being the underlying mechanism for growth with Gd^3+^.

**FIG 1.**
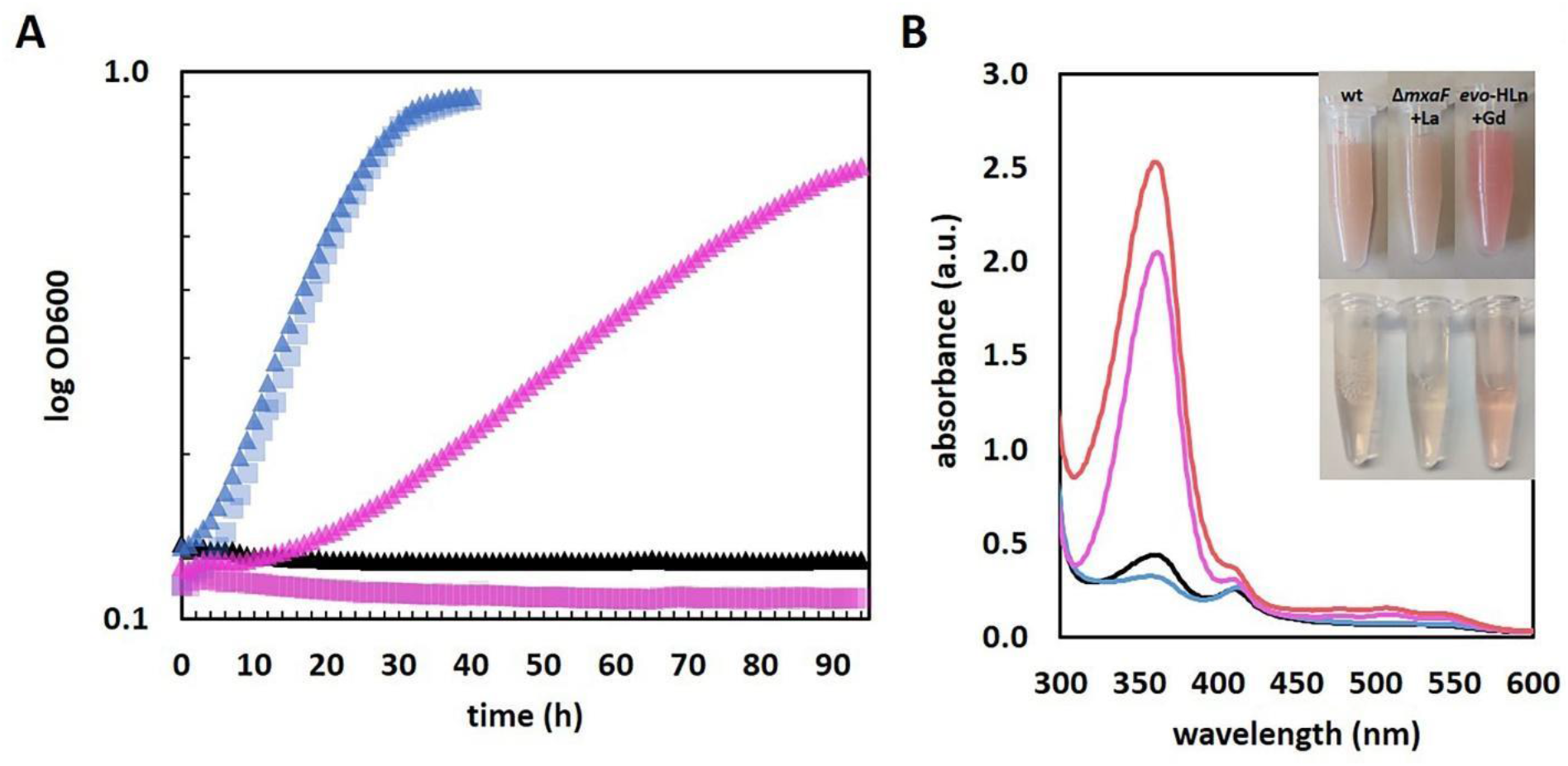
Growth of *M. extorquens* AM1 *evo*-HLn with heavy Ln^3+^. *A*, Δ*mxaF* (squares) and *evo*-HLn (triangles) strains were grown in methanol minimal medium with either no Ln^3+^ (black), 2 μM Gd^3+^ (pink) or 2 μM La^3+^ (blue). Not all data points are visible for strains and conditions with no growth. *B*, UV-visible spectrum of cell extracts of wild type grown without Ln^3+^ (black), *evo*-HLn with 2 μM Gd^3+^ (pink), and Δ*mxaF* with 2 μM La^3+^ (blue). Absorbance at 360 nm increases in the *evo*-HLn extract with the addition of 130 μM PQQ (red). Cells extracts were prepared in 25 mM Tris, pH 8.0. Inset shows visual color differences among strains and conditions. Spectra represent the average of 3 separate replicates with extracts containing 5.3-5.6 mg/mL protein. *Inset top*, cell suspension; *inset bottom*, cell extract.

Genomic DNA was isolated from the variant, sequenced, and analyzed for mutations relative to ancestral Δ*mxaF* strain. Three single nucleotide polymorphisms (SNPs) were identified in the variant compared to Δ*mxaF* (Table S1). Only one of the three mutations was categorized as non-synonymous: a T to A nucleotide transition, resulting in a leucine to histidine amino acid substitution in a hybrid histidine kinase/response regulator (META1_1800). The mutation was confirmed by Sanger sequencing analysis, and the variant strain was named *evo*-HLn for “evolved for growth with heavy lanthanides”. To determine SNP conferred a gain-of-function mutation, the mutant allele was cloned into an IPTG-inducible expression vector, which was then transformed into the Δ*mxaF* ΔMETA1_1800 double knockout mutant strain. When expression of the mutant allele was induced, we observed Gd^3+^-dependent methanol growth (final OD = 0.5; N = 4). No growth was observed for the uninduced control condition.

### Increased PQQ biosynthesis

We observed that the cells of *evo*-HLn grown in methanol minimal medium with Gd^3+^ had a distinctive, bright pink coloration, and that extracts prepared from *evo*-HLn cells retained this increased pigmentation (Fig. 1B inset). When analyzed by UV-visible spectrophotometry, *evo*-HLn extracts displayed a unique peak at 361 nm (Fig. 1B). A peak around this wavelength is a signature of PQQ when bound to XoxF MDH or ExaF EtDH (Good et al. 2016, 2019). To confirm PQQ was the cause of the absorption anomaly, we spiked it into the *evo*-HLn extracts and observed an increase at the same wavelength. After normalizing for protein concentrations, the absorbance spectra indicated PQQ in *evo*-HLn extracts was 4-fold higher compared to wild type and 6-fold higher compared to Δ*mxaF* extracts.

### Increased *xox1* promoter and MDH activities

Since XoxF MDH is closely linked with Ln^3+^-dependent methanol growth, one plausible explanation for the expanded range of metals used by *evo*-HLn was increased XoxF MDH activity. Reporter-fusion assays previously showed that *xox1* promoter activity was stimulated by light Ln^3+^ ranging from La^3+^ to neodymium (atomic number 60), with only a minor increase above background activity with Sm^3+^ (Vu et al. 2016). We measured *xox1* promoter activity in *evo*-HLn with La^3+^ and observed a 7-fold increase compared to Δ*mxaF* and an 11-fold increase compared to wild type (Fig. 2A). Next, we measured *xox1* promoter activity with Gd^3+^ from *evo*-HLn and observed a similar increase. Further, we did not detect *xox1* promoter activity in wild type with Gd^3+^ (Fig. 2A), showing that although the wild type grows with methanol in the presence of Gd^3+^ (Fig. S2), the regulatory switch from MxaFI MDH to XoxF MDH oxidation systems does not occur. This could be indicative of either the wild type being unable to transport Gd^3+^ or Gd^3+^ not functioning as a signal for the “lanthanide switch” in this strain. Regardless, it can be concluded that wild type grows on methanol using MxaFI MDH, the Ca^2+^/PQQ-dependent oxidation system, when Gd^3+^ is present in the medium.

**FIG 2.**
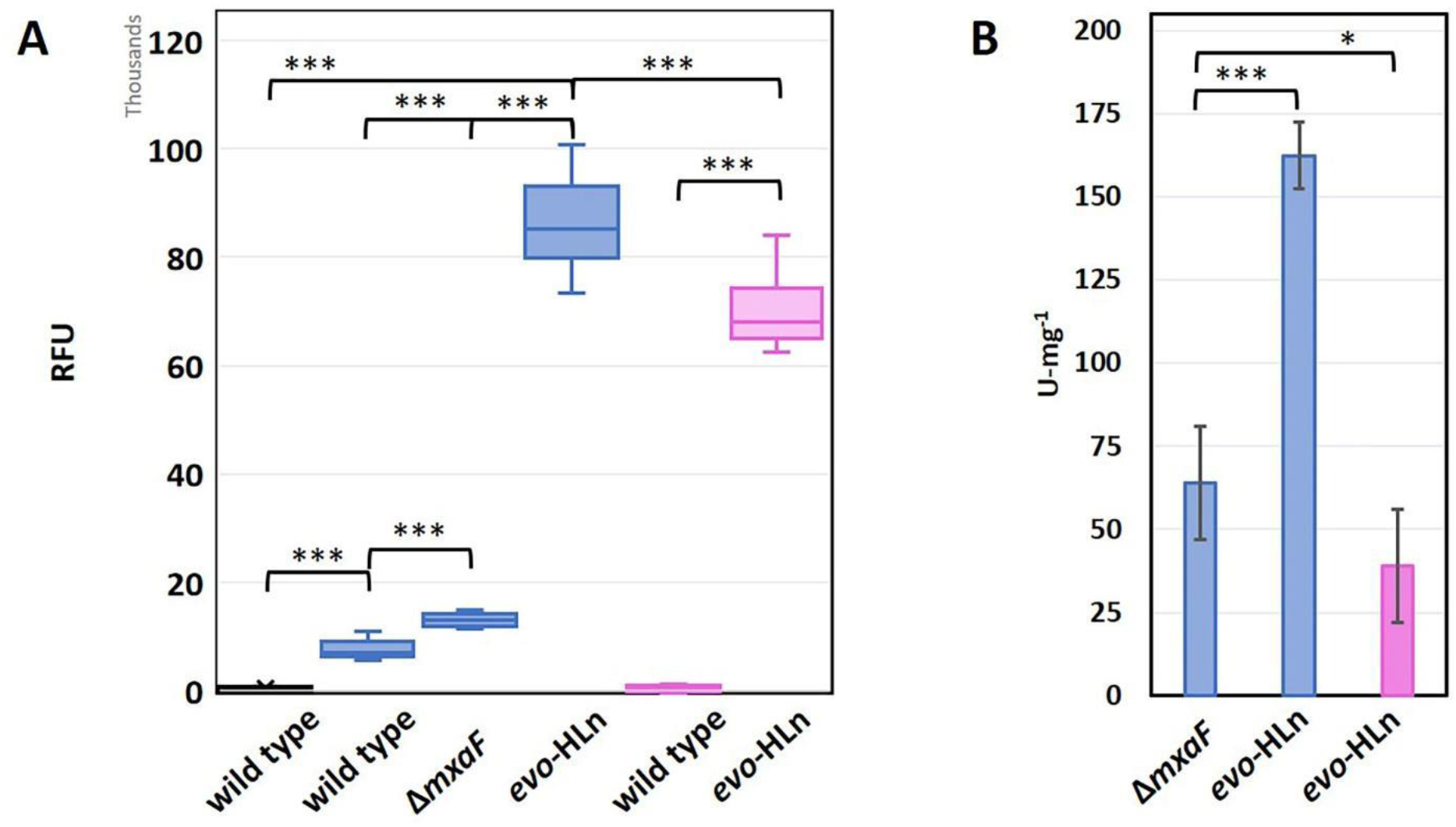
*evo*-HLn exhibits higher *xox1* promoter and Ln^3+^-dependent MDH activities with both light and heavy Ln^3+^. *A*, wild type, Δ*mxaF* and *evo*-HLn carrying a *xox1 promoter-yfp* reporter fusion construct were grown with methanol and either no Ln^3+^ (black), La^3+^ (blue), or Gd^3+^ (pink) to an OD of ~0.4 at 600 nm and promoter readout was measured as fluorescence. Box and whisker plot shows the interquartile range of RFU determined for 9-12 biological replicates from 3 independent experiments. Whiskers show the minimum and maximum values. Median values marked by a bold line, and the mean as an X. For each strain and growth condition, readout from the promoter-less construct was subtracted as background fluorescence. *B*, MDH activity measured from cell extracts of Δ*mxaF* and *evo*-HLn grown in methanol medium with either La^3+^ (blue), or Gd^3+^ (pink). MDH activity was determined using the DCPIP dye-linked assay according to Anthony and Zatman with previously reported modifications (25, 67). For both panels *A* and *B*, *** is significant at *p* < 0.001 and * at *p* < 0.05 by One-Way ANOVA and Tukey’s Honestly Significant Difference (HSD) test.

Next, we measured MDH activity in cell-free extracts of Δ*mxaF* and *evo*-HLn prepared from cultures grown with methanol and either La^3+^ or Gd^3+^. When grown with La^3+^, MDH activity in *evo*-HLn extracts was ~3-fold higher than in Δ*mxaF* extracts (Fig. 2B), verifying increased production of XoxF enzyme. Ln^3+^ species do not function equally well as part of the XoxF MDH cofactor complex, along with PQQ, and the enzyme active site is finely tuned for light Ln^3+^ (Jahn et al. 2018; Daumann 2019). Therefore, a reduction in XoxF MDH function could be expected with Gd^3+^ in the active site. MDH activity was detectable in extracts of *evo*-HLn grown with Gd^3+^ corresponding to 68% of the activity measured in extracts of Δ*mxaF* with La^3+^ (Fig. 2B). *evo*-HLn grows well with Gd^3+^, and increased production of XoxF MDH is likely a major contributor to this metabolic capability. Increased *xox1* promoter and MDH activities of *evo*-HLn are suggestive of possible increases in Ln^3+^ transport and intracellular accumulation.

### Enhanced Lanthanide accumulation in *evo*-HLn

Using inductively-coupled plasma mass spectroscopy (ICP-MS), we determined the Ln^3+^ metal content of cells grown with methanol and a single Ln^3+^ element species. Uptake and storage of Gd^3+^ by *evo*-HLn was a striking ~5 mg/g CDW. Wild type Gd^3+^ content was ~36-fold less, but it was measurable (Fig. 3A). Transport and storage of Gd^3+^ by the wild type was somewhat surprising as the strain does not grow on methanol if it is the only Ln^3+^ provided to the growth medium. The opposite trend, though much less pronounced, was observed when measuring accumulation of La^3+^ with *evo*-HLn accumulating ~3-fold less than the wild type. Comparing uptake and storage of each Ln^3+^ within the same strain, we observed that *evo*-HLn accumulated ~82-fold more Gd^3+^ than La^3+^. The wild type, on the other hand, accumulated ~43% more La^3+^ than Gd^3+^. These data suggested that the wild type prefers uptake and storage of light Ln^3+^, such as La^3+^, over heavy Ln^3+^. In contrast, increased Gd^3+^ accumulation seen for *evo*-HLn suggested that it may have evolved a preference for heavy Ln^3+^. To test this possibility, we next compared Ln^3+^ accumulation when the strains were provided equal concentrations of both La^3+^ and Gd^3+^ in the growth medium. Wild type accumulated equal amounts of La^3+^ and Gd^3+^ (Fig. 3B). Total Ln^3+^ content was ~1.7-fold the amount of La^3+^ and ~2-fold the amount of Gd^3+^ that was stored when only the single Ln^3+^ species was provided (Fig. 3AB). Accumulation of the individual Ln^3+^ species by *evo*-HLn was also equal when both were provided, but the overall levels were ~3-fold less than what was observed for the wild type (Fig. 3B). Total Ln^3+^ content was ~1.5-fold that of La^3+^ alone for *evo*-HLn, but ~63-fold less than the amount of Gd^3+^ alone (Fig. 3AB). These data suggest that light Ln^3+^ may impact the capacity for *evo*-HLn to acquire and store heavy Ln^3+^ and may be an indication of a second transport system for the latter.

**FIG 3.**
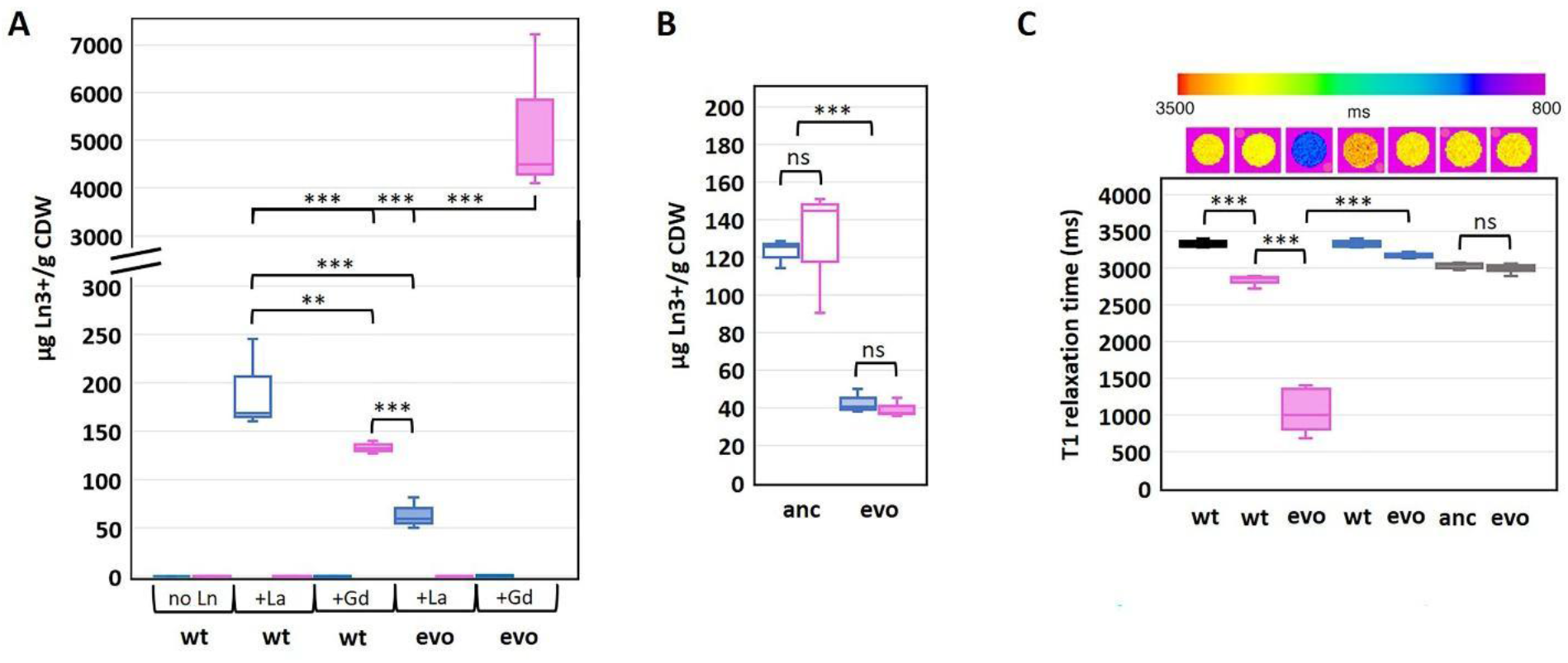
Ln^3+^ accumulation during methanol growth. *A*, Intracellular Ln^3+^ content in *evo*-HLn and wild type grown on methanol minimal medium with 2 μM GdCl_3_ 2 μM LaCl_3_ or no lanthanides (Ln) as indicated. Gd^3+^ (pink) and La^3+^ (blue) contents are normalized to cell dry weight (CDW) for each growth condition. Box plots show interquartile range (boxes), median (line), and standard deviation (bars) for three biological replicates, each quantified three times by ICP-MS. *B*, Intracellular Ln^3+^ concentrations in the ancestral Δ*mxaF* mutant (open boxes) and *evo*-HLn (shaded boxes) when cells were grown with 2 μM GdCl_3_ and 2 μM LaCl_3_. Plots represent data processed the same way as in *A*. *C*, MRI T1 relaxation time measurements of whole cells grown with Ln^3+^. Top shows digital scan with T1 times color coded. Bottom shows T1 relaxation times of cells grown with La^3+^ (blue), Gd^3+^ (pink), or both (black). Boxes represent the upper and lower quartiles of three independent biological replicate samples. Median lines and standard deviations (bars) are shown. ** is significant at *p* value < 0.01 and *** is significant at *p* value < 0.00001 by One-Way ANOVA and Tukey’s HSD test. ns is not statistically significant at *p* < 0.05. For *A, B*, and *C*: wt, wild type; evo, *evo*-HLn; anc, Δ*mxaF*.

### Gd^3+^ accumulation in *evo*-HLn shortens MRI T1 relaxation time in whole cells

Since *evo*-HLn was able to transport and accumulate increased levels of Gd^3+^, we tested if the paramagnetic metal could affect MRI contrast. When we scanned whole cells by MRI, we observed a statistically significant reduction in T1 relaxation time for cells grown with Gd^3+^ compared to cells grown with La^3+^ or without Ln^3+^ (Fig. 3C). *evo*-HLn cells cultured with Gd^3+^ displayed a 3-fold decrease in T1 relaxation time when compared to wild type cultured without Ln^3+^. The T1 relaxation time of wild type cells with Gd^3+^ was 17% less than cells without Ln^3+^. In contrast, intracellular accumulation of La^3+^ by cells had very little impact on T1 relaxation times (Fig. 3C). When *evo*-HLn and Δ*mxaF* cells were grown with both Gd^3+^ and La^3+^, T1 relaxation times were shortened by ~10% compared to the no Ln^3+^ treatment. These data show that MRI is sensitive enough to detect Gd^3+^ accumulation in *M. extorquens* AM1 and that *evo*-HLn cells can accumulate Gd^3+^ to intracellular concentrations that are high enough to produce robust MRI contrast.

### Efficient acquisition of Gd^3+^ from the GBCA Gd-DTPA

Finally, the capacity of the *evo*-HLn strain to acquire Gd^3+^ from the chelator diethylenetriamine pentaacetate (DTPA) was demonstrated (Fig. 4). Despite the high stability of the Gd-DTPA complex (log *K*_therm_ 22, log *K*_cond_ 17; (Tweedle et al. 1991; Wedeking, Kumar, and Tweedle 1992)), *evo*-HLn was able to grow readily with no reduction growth rate compared to growth with soluble GdCl_3_ (Gd-DTPA, 0.04 h^-1^ ± 0.00; GdCl_3_, 0.03 h^-1^ ± 0.00; n = 3). This result indicates that *evo*-HLn has a highly effective means of sequestering Gd^3+^ from DTPA, thus demonstrating a potential importance as a key player in Gd^3+^ recycling and pollution remediation.

**FIG 4.**
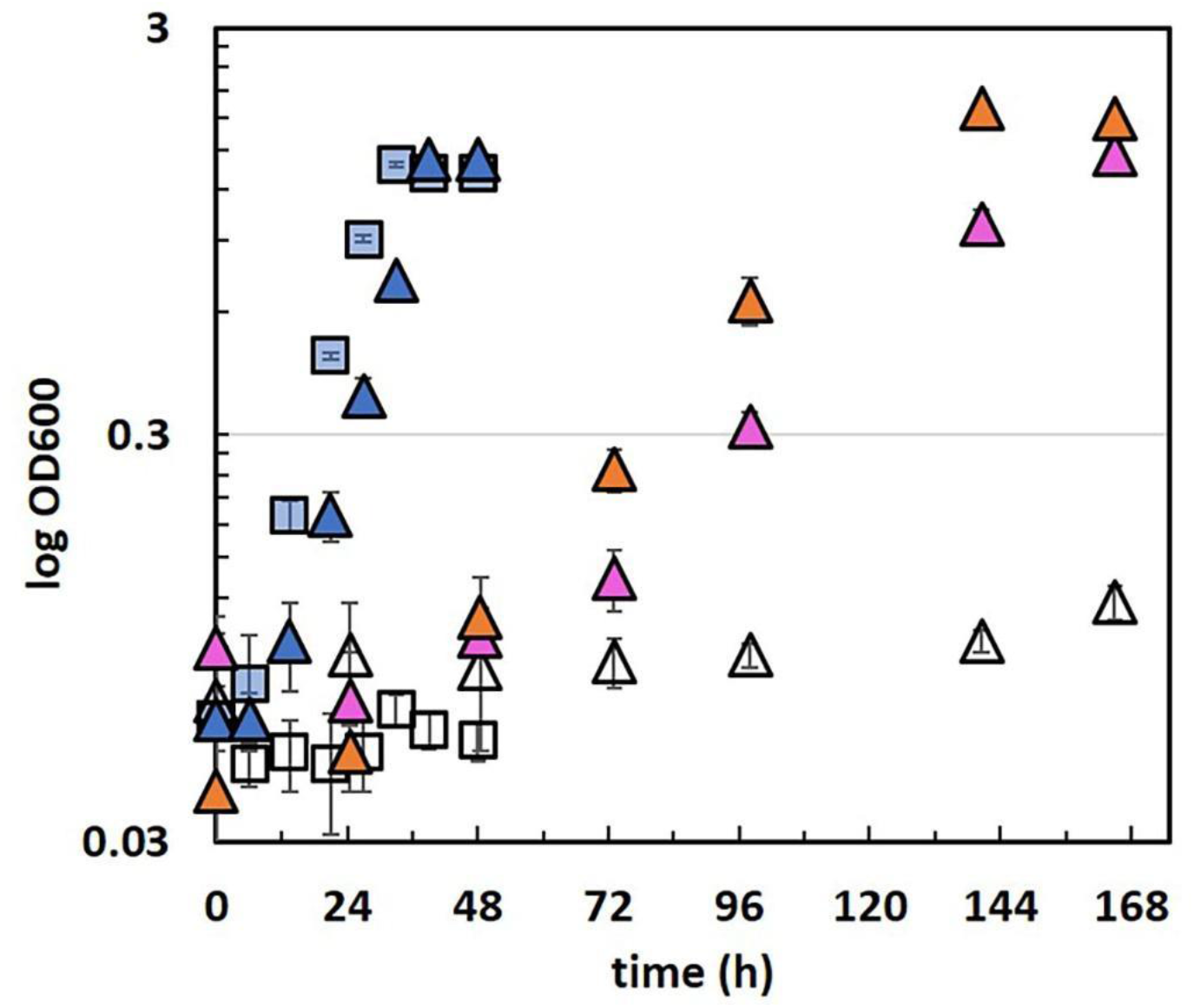
Methanol growth of *evo*-HLn with the GBCA Gd-DTPA. Density of *evo*-HLn cultures is shown as triangles. Ln^3+^ sources are denoted by color. Pink, 2 μM GdCl_3_; orange, 2 μM Gd-DTPA; open symbols, no Ln; blue, 2 μM LaCl_3_. The ancestral strain, Δ*mxaF* (squares), grown with 2 μM La^3+^ is included for comparison. Data points represent the average of 6 biological replicates from 2 independent experiments with error bars showing standard deviations.

## DISCUSSION

It is common practice to inject patients with GBCAs before MRI to enhance scan effectiveness by shortening the T1 relaxation of the target. Chelate composition and complexation can significantly affect both T1 relaxation and the stability of GBCA complexes (Caravan 2006). Moreover, Gd^3+^ is conjugated to proteins for applications such as blood pool agents, which can be done chemically (Caravan 2006) via avidin-biotin (Dafni et al. 2002; Gilad et al. 2005) and genetically, via lanthanide binding tags (Xue et al. 2015). Alternatively, Gd^3+^ GBCAs are used as substrates for genetically encoded reporters (Nyström et al. 2019). Therefore, identifying novel peptides and proteins that effectively chelate Gd^3+^ will aid in the development of next-generation GBCAs. We show that the genetic variant of *M. extorquens* AM1, *evo*-HLn, has acquired the ability to transport and accumulate Gd^3+^ such that a significant contrast can be observed by MRI. These results indicate *evo*-HLn produces proteins and/or peptides for these physiological processes that efficiently bind free Gd^3+^. *M. extorquens* AM1 produces several Ln^3+^-binding molecules for uptake and utilization encoded in the *lut* gene cluster (Roszczenko-Jasińska et al. 2020). Several lines of evidence also support the existence of excreted molecules for extracellular Ln^3+^ acquisition, called lanthanophores (Ochsner et al. 2019; Roszczenko-Jasińska et al. 2020; Daumann 2019). It is likely that *evo*-HLn uses some or all this uptake and utilization machinery to grow with Gd^3+^, but the Ln^3+^ accumulation data reported here suggests there may be alternative or additional machinery for uptake of heavy Ln^3+^. Though the wild type can transport and accumulate Gd^3+^ intracellularly, it is not sufficient to function as a signal for the switch from MxaFI MDH to XoxF MDH. These observations are suggestive of additional components, including regulatory elements, being utilized or co-opted for Gd^3+^ acquisition, uptake, transport and/or utilization in *evo*-HLn. Identification of these additional peptides/proteins may provide new candidate chelators or scaffolds for protein engineering of more bio-safe chelates.

Although Gd^3+^ is considered safe when properly chelated, the thermodynamic and kinetic stabilities of GBCAs differ depending on the chemical structure (Sherry, Caravan, and Lenkinski 2009). Deposition of Gd^3+^ correlates with detrimental health effects in humans, but long-term environmental studies investigating ecotoxicological effects and bioaccumulation have yet to be pursued (Ebrahimi and Barbieri 2019; Thomsen 2017). We show that *evo*-HLn grows readily on methanol with Gd-DTPA as the sole Ln^3+^ source, revealing that *M. extorquens* AM1 must produce molecular machinery capable of sequestering Ln^3+^ from Gd-complexes. It’s been speculated that the Ln^3+^-binding peptide lanmodulin may be important for transport of light Ln^3+^ (Cotruvo et al. 2018), but it has been shown that a null mutation in the gene encoding this peptide does not impact La^3+^-dependent methanol growth in *M. extorquens* PA1 (Ochsner et al. 2019), a closely-related strain, nor was the gene hit in a mutant hunt to detect essential genes for La^3+^-dependent methanol oxidation in *M. extorquens* AM1 (Roszczenko-Jasińska et al. 2020). Further studies are needed to define if lanmodulin is necessary for Gd^3+^ transport or if an alternative system is necessary for this process. In addition, we provide evidence that *evo*-HLn transports and stores more Gd^3+^ than wild type does La^3+^ or Gd^3+^. The capability for growth with GBCAs and increased uptake and storage of Gd^3+^ make *evo*-HLn an excellent candidate for the development of a microbial platform for recovery of Gd^3+^ from wastewater and possibly even medical waste.

The implications of *evo*-HLn on our understanding of Ln^3+^-biochemistry are extensive. First, *evo*-HLn is the first genetic variant reported to be adapted for growth with heavy Ln^3+^. Our results indicate wild type is unable to efficiently transport Gd^3+^, raising the question of exactly how *evo*-HLn has adapted to efficiently acquire, transport and use the heavy Ln^3+^ (Fig. 5). Regarding acquisition, it is possible that *evo*-HLn produces a modified lanthanophore or a novel Ln^3+-^-chelating molecule. We showed that *evo*-HLn, when grown with the heavy lanthanide Gd^3+^, produces higher levels of PQQ. This is striking as PQQ biosynthesis genes have been reported to be down-regulated with the light lanthanide La^3+^ (Good et al. 2019). This is supported by the drop of absorbance at 360 nm seen in the Δ*mxaF* extracts compared to the wild type. PQQ, already shown to bind Ln^3+^ in solution (Lumpe and Daumann 2019), may therefore be critical for binding heavy Ln^3+^ and/or could serve as a signal molecule for their uptake. Increased *xox1* promoter and XoxF1 MDH activities reflect regulatory changes necessary for methanol growth with Gd^3+^. However, this regulatory change appears to be insensitive to light versus heavy Ln^3+^ distinction, as shown by augmented XoxF1 activity with La^3+^. In addition, increased Ln^3+^ transport suggests disruption of the tightly controlled regulatory mechanisms governing uptake (Roszczenko-Jasińska et al. 2020). The details of the molecular process and regulation of storage are still unknown, but increased Ln^3+^ accumulation in *evo*-HLn provides an excellent comparator to wild type and ancestral strains to investigate these questions in depth.

**FIG 5.**
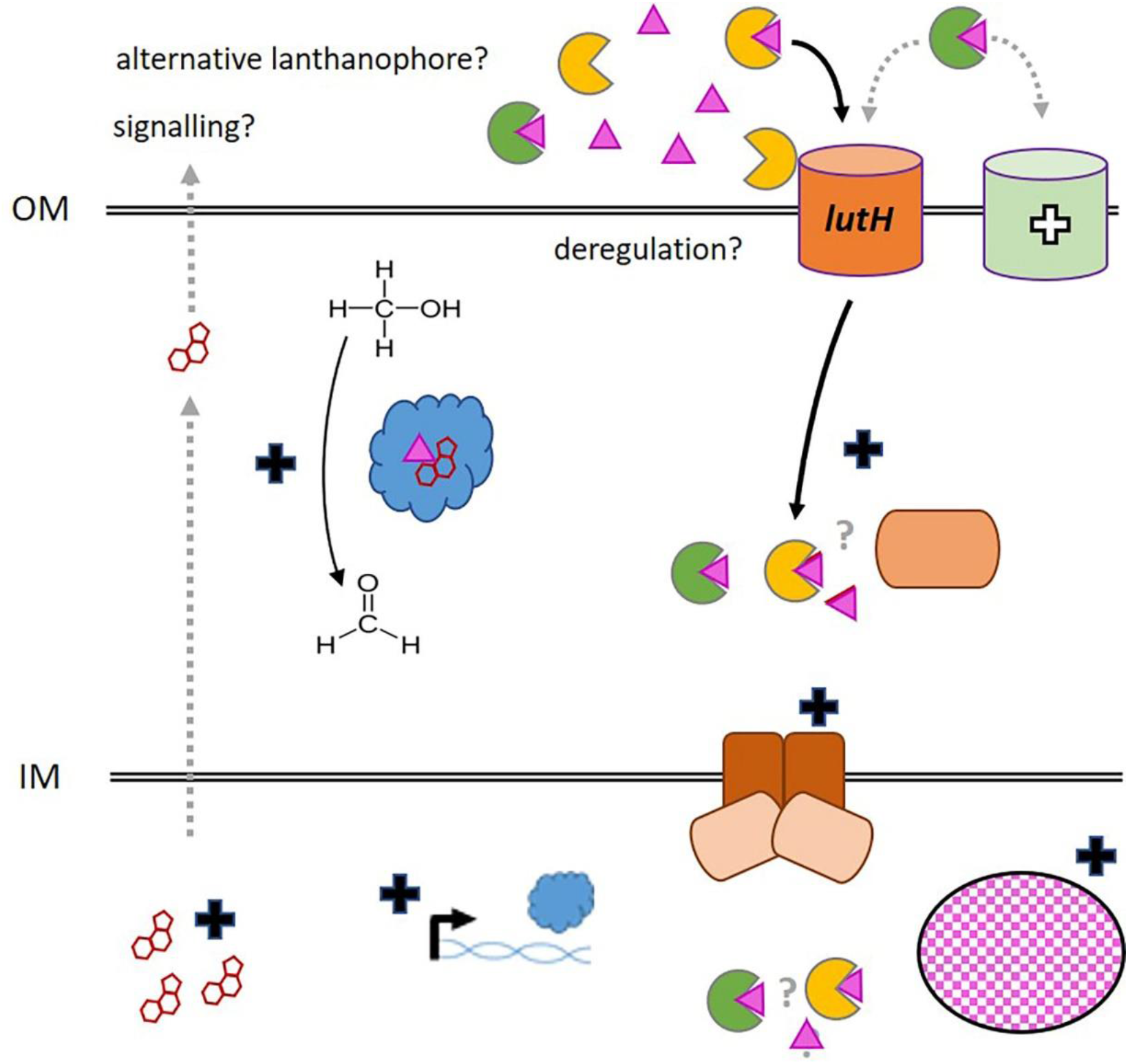
Physiological implications of genetic adaptation for growth with heavy Ln^3+^ in *evo*-HLn. Crosses indicate a change in activity or function relative to the ancestral Δ*mxaF* strain: white, novel; black, increased. Exogenous Gd^3+^ (pink triangles) is acquired by excreted metal-chelating molecules, called lanthanophores (yellow notched circles). Uptake of Gd^3+^ may involve an alternative lanthanophore (green notched circle). Increased Ln^3+^ transport suggests possible disruption of the tight regulation of *lut* gene expression (gene products shown in orange, TonB-dependent receptor, ABC transport periplasmic, membrane, and cytoplasmic components) or upregulation of an alternative transport system for heavy Ln. Augmented Gd^3+^ storage is denoted as the pink checked oval. Increased *xoxF1* expression and XoxF1 MDH activity is shown in blue. PQQ biosynthesis is also elevated. Gray dashed arrow represents a possible role in signalling or metal chelation for PQQ (red rings). Gray question marks signify that the details of the state of Ln^3+^ in the periplasm for transport to the cytoplasm, whether free or complexed with a lanthanophore, are not yet resolved. OM, outer membrane; IM, cytoplasmic membrane.

Together, the observations reported here indicate changes in numerous physiological processes in *evo*-HLn allowing for growth with heavy Ln^3+^. RNA-sequencing of *evo*-HLn growth with Gd^3+^, which is already underway, will give invaluable insight into the regulatory change(s) underlying these physiological alterations. Unraveling the links between genetic adaptation(s) and physiological changes will further illuminate our understanding of Ln^3+^ as life metals as well as aid in the development of microbial technologies for Gd^3+^ bioremediation and contrast agents.

## MATERIALS AND METHODS

### Strains and culture conditions

*M. extorquens* AM1 strains were routinely grown at 30°C MP minimal medium (Delaney et al. 2013) with 15 mM succinate, shaking at 200 rpm on an Innova 2300 platform shaker (Eppendorf, Hamburg, Germany). For growth studies, 50 mM methanol was used as the sole carbon and energy source. Lanthanides were added as chloride salts or gadopentetic acid (Gd-DTPA; Magnevist^®^) to a working concentration of 2 or 20 μM as indicated. When necessary, 50 μg/mL kanamycin was added to the growth medium for plasmid maintenance. Strains and plasmids used in this study are listed in Table S1 of the supplementary material.

### Plasmid construction

The META1_1800 mutant allele was synthesized as a blunt-end 5’-phosphrylated 1436 bp gBlock (IDT, Coralville, IA, USA) with the modified *lac* promoter P*_L_*/O4/A1 (Carrillo et al. 2019) upstream. The gBlock was ligated into pCM66T digested with Ecl136II (Thermo Fisher Scientific, Waltham, MA, USA) to generate pNG327.

### Strain construction

*M. extorquens* AM1 strains were transformed by electroporation (Toyama, Anthony, and Lidstrom 2006). After 24 hours of outgrowth, transformants were selected by plating on MP medium with 1.5% agar, 15 mM succinate and 50 μg/mL kanamycin or 20 μg/mL tetracycline. Transformants were incubated at 30 °C until isolated colonies appeared.

### Methanol growth analysis with light and heavy lanthanides

*M. extorquens* AM1 strains were grown with succinate overnight, cells were pelleted by centrifugation at 1,000 x g for 10 min at room temperature using a Sorvall Legend X1R centrifuge (Thermo Scientific, Waltham, MA, USA), and washed in 1 mL of sterile MP medium with methanol. For growth analysis in microplates, washed cells were resuspended in 200 μL of MP methanol medium and 10 μL were transferred to each microplate well with 640 μL MP methanol medium. For growth studies with Gd-DTPA, 50 μL of inoculum was added to 3 mL MP methanol medium in sterile 14 mL polypropylene culture tubes (Fisher Scientific, Hampton, NH, USA). Culture densities were monitored over time by measuring light scatter at 600 nm using either a Synergy HTX multi-mode plate reader (Biotek, Winooski, VT, USA) or an Ultraspec 10 density meter (Biochom, Holliston, MA, USA).

### UV-visible spectrophotometry

To prepare cell-free extracts, 50 mL of methanol grown culture with Gd^3+^ or La^3+^ was harvested, upon reaching an OD_600_ of ~1.1-1.3, by centrifugation at 4,696 x *g* for 10 minutes at 4 °C. The supernatant was removed and cell pellets were resuspended in 1.5 mL of 25 mM Tris, pH 8.0 and lysed using an OS Cell Disrupter at 25,000 psi (Constant Systems Limited, Low March, Daventry, Northants, United Kingdom). Lysates were transferred to 1.5 mL eppendorf tubes and clarified of cell debris by centrifugation at 21,000 x *g* for 10 minutes at 4 °C. Cell-free extracts were transferred to new eppendorf tubes and kept on ice until needed. PQQ was prepared fresh to a working concentration of 5.3 mM in an opaque conical tube and kept on ice until needed. Absorbance spectra were measured from 250-600 nm with a Synergy HTX multi-mode plate reader. A blank buffer spectrum was subtracted as background. Protein concentrations were determined by absorbance at 280 nm and the bicinchoninic acid assay (ThermoFisher Scientific, Waltham, MA, USA).

### Genomic DNA extraction and sequencing

The Δ*mxaF* and *evo*-HLn mutant strains were grown in shake flasks with 50 mL MP with succinate to OD_600_ ~ 1.0. Genomic DNA was extracted according to the “Bacterial genomic DNA isolation using CTAB” protocol (Joint Genome Institute, Walnut Creek, CA, USA). Samples were submitted to Genewiz (South Plainfield, NJ, USA) for whole genome sequencing using the Illumina HiSeq platform with 2 x 150 bp read length. Variant calling and analysis was performed by Genewiz.

### Transcriptional reporter fusion assays

Strains carrying VENUS *yfp* fusion constructs were grown on methanol in 48-well microplate format. Upon reaching a culture density of OD_600_ ~0.35, 200 μL of culture were transferred to an optical bottom black 96-well plate. Fluorescence was measured at an excitation wavelength of 485 nm and an emission wavelength of 520 nm. Relative fluorescence units (RFU) were calculated as raw fluorescence (F_520_ nm) divided by OD_600_.

### Methanol dehydrogenase activity assays

Cell extracts were prepared as described above, but with an additional wash step in 20 mL of 100 mM Tris-HCl, pH 9.0 before lysing. Protein concentrations of cell-free extracts were determined by BCA assay. Methanol dehydrogenase activity was measured by monitoring the phenazine methosulfate (PMS)-mediated reduction of 2,6-dichlorophenol indophenol (DCPIP; ε_600nm_ = 21 mM^-1^ cm^-1^ (Good et al. 2016, 2019, 2020)) as described (C. Anthony and Zatman 1967; R. Ghosh and Quayle 1979; Vu et al. 2016; Good et al. 2020). To reduce background activity, all assay reagents were dissolved in water; PES and DCPIP solutions were prepared in opaque tubes and kept on ice; and cell-free extracts were pre-incubated for 2 minutes at 30 °C as recommended (Jahn et al. 2020).

### Whole-cell MRI

Wild type and *evo*-HLn mutant strains were grown in 50 mL methanol medium to maximal culture density (OD_600_ ~3). No REE was added to the medium for the wild-type strain. Either Gd^3+^, La^3+^, or both (2 μM each) was added to the medium for *evo*-HLn. Cells were harvested by centrifugation at 4,696 x *g* for 10 minutes at room temperature. The supernatant was removed, cells were washed two times by resuspension with 50 mL of 25 mM Tris, pH 7.0, and centrifugation at 4,696 x *g* for 10 minutes at room temperature. After washing, 1/10 of the cell pellets were resuspended in 500 μL of 25 mM Tris, pH7.0. MRI data was acquired using a 7.0 T horizontal MRI (Bruker) equipped with a multi-channel receive array and volumetric transmit coil ensemble. T1 maps were generated via Paravision 360, with 10 TRs ranging from 400-17500, TE = 6.89, MTX = 128/128, FOV = 32cm^2^, ST = 1mm and AVG = 3.

### Intracellular Ln^3+^ quantification

After whole-cell MRI analysis, cell pellets were dehydrated at 65 °C for 72 hours. Dried pellets were weighed before deconstruction in *Aqua regia* diluted in 2% nitric acid and sonicated for 0.5 h before passing through 0.45 μm Whatman syringe filters. Samples were sent to the Laboratory for Environmental Analysis (Center of Applied Isotope Studies, University of Georgia) for ICP-MS analysis.

## ACKNOWLEDGMENTS

We thank Elizabeth Skovran for kindly providing us pAP05 and pES508. We thank Lena Daumann for generously providing PQQ powder, and for her valuable suggestions on this manuscript.

This material is based upon work supported by the National Science Foundation under Grant No. 1750003 for N.C.M.G. and N.M.G. We acknowledge funding for A.A.G. from NIH/NINDS: R01-NS098231/R01-NS104306 NIH/NIBIB: P41-EB024495; NSF 2027113.

Conceptualization, N.M.G., N.C.M.G., and A.A.G.; methodology, N.M.G.; investigation, N.M.G., H.L., E.R.H.; writing - original draft, N.M.G., H.L.; writing - reviewing and editing, all authors; funding acquisition, N.C.M.G. and A.A.G.; resources, N.C.M.G. an A.A.G.

N. C. M. G. and N. M. G are inventors on a patent application submitted by the Regents of the University of California

## TABLES

**TABLE 1.** Growth rates and yields of strains grown in minimal medium with methanol Ln^3+^. Culture density was monitored for up to 96 hours.

| strain | $\text{Ln}^{3+}$ source<sup>ε</sup> | growth rate<sup>9</sup> | growth yield<sup>9</sup> |
| --- | --- | --- | --- |
| wild type | none | $0.15 \pm 0.01$ | $0.72 \pm 0.13$ |
| wild type | $\text{LaCl}_3$ | $0.15 \pm 0.01$ | $0.78 \pm 0.14$ |
| wild type | $\text{GdCl}_3$ | $0.14 \pm 0.00$ | $0.77 \pm 0.14$ |
| $\Delta mxaF$ | none | n.d. | - |
| $\Delta mxaF$ | $\text{LaCl}_3$ | $0.14 \pm 0.01$ | $0.90 \pm 0.04$ |
| $\Delta mxaF$ | $\text{GdCl}_3$ | n.d. | - |
| *evo*-HLn | none | n.d. | - |
| *evo*-HLn | $\text{LaCl}_3$ | $0.11 \pm 0.01$ | $0.93 \pm 0.13$ |
| *evo*-HLn | $\text{GdCl}_3$ | $0.03 \pm 0.00$ | $0.69 \pm 0.04$ |

<sup>ε</sup> $\text{Ln}^{3+}$ were provided in the growth medium at 2 $\mu\text{M}$
<sup>9</sup> Values represent the averages of 10 biological replicates from 3 independent experiments except where indicated. Error bars are standard errors of the mean (SEM). n.d. is not determined. - is no growth.

