## Supplemental Tables and Figures for "Harnessing methylotrophs as a bacterial platform to reduce adverse effects of the use of the heavy lanthanide gadolinium in magnetic resonance imaging"

**SUPPLEMENTARY MATERIAL**

**TABLES**

**TABLE S1. Bacterial strains and plasmids used in this study**

| strain or plasmid | description | reference |
|---|---|---|
| **strains** | | |
| ***Methylorubrum extorquens*** | | |
| AM1 | wild type; rifamycin-resistant derivative | (1) |
| Δ*mxaF* | deletion mutant | (2) |
| *evo*-HLn | Δ*mxaF* deletion mutant variant adapted for methanol growth with heavy lanthanides | this study |
| **plasmids** | | |
| pNG327 | $P_L$/O4/A1 expression vector with *evo*-HLn variant META1_1800 allele, $Km^r$ | this study |
| pAP05 | promoterless *yfp* fusion vector, $Tc^r$ | (3) |
| pES503 | pAP05 with *xox1* promoter region, $Tc^r$ | (3) |

**TABLE S2. Mutations detected by genome resequencing of Δ*mxaF* and *evo*-HLn**

Wild-type *M. extorquens* AM1 was used as the reference strain for mapping. Green, mutations unique to Δ*mxaF*; yellow, mutations unique to*evo*-HLn.

| Chromosome | Region | Type | Ref | Allele | Count | Freq | Qual | locus_tag | Coding change | Amino acid change | Non |
|---|---|---|---|---|---|---|---|---|---|---|---|
| CP001510 | 482893^482894 | In | - | C | 47 | 100 | 200 | META1p0458 | | | - |
| CP001510 | 1673173 | SNV | A | G | 110 | 100 | 200 | META1p1592 | 69T>C | | No |
| CP001510 | 1873778 | SNV | T | A | 175 | 100 | 200 | META1p1800 | 452T>A | Leu151His | Yes |
| CP001510 | 2329711 | Del | G | - | 125 | 98 | 160 | | | | - |
| CP001510 | 2777457 | SNV | T | G | 38 | 97 | 160 | META1p2648 | 408A>C | | No |
| CP001510 | 2803789 | SNV | C | T | 32 | 100 | 200 | META1p2676 | .63C>T | | No |
| CP001510 | 2803840 | SNV | T | C | 9 | 100 | 155 | META1p2676 | 114T>C | | No |
| CP001510 | 2891642 | SNV | G | C | 185 | 100 | 200 | META1p2763 | 879C>G | | No |
| CP001510 | 3037769 | Del | C | - | 190 | 95 | 200 | META1p2908 | 718delC | Arg241fs | Yes |
| CP001510 | 3159071 | Del | G | - | 94 | 97 | 160 | | | | - |
| CP001510 | 4001527..4001531 | Del | CGTGC | - | 122 | 85 | 200 | META1p3891, META1p3892 | 262_266delGCACG | Ala88fs | Yes |
| CP001510 | 4322600 | SNV | G | A | 4 | 67 | 26 | META1p4234 | 960C>T | | No |
| CP001511 | 580985^580986 | In | - | C | 128 | 98 | 200 | META2p0619 | 468_469insG | Arg157fs | Yes |
| CP001511 | 770863 | Del | G | - | 52 | 95 | 160 | META2p0816 | 890delG | Ala298fs | Yes |

**FIGURES**

**FIG S1**

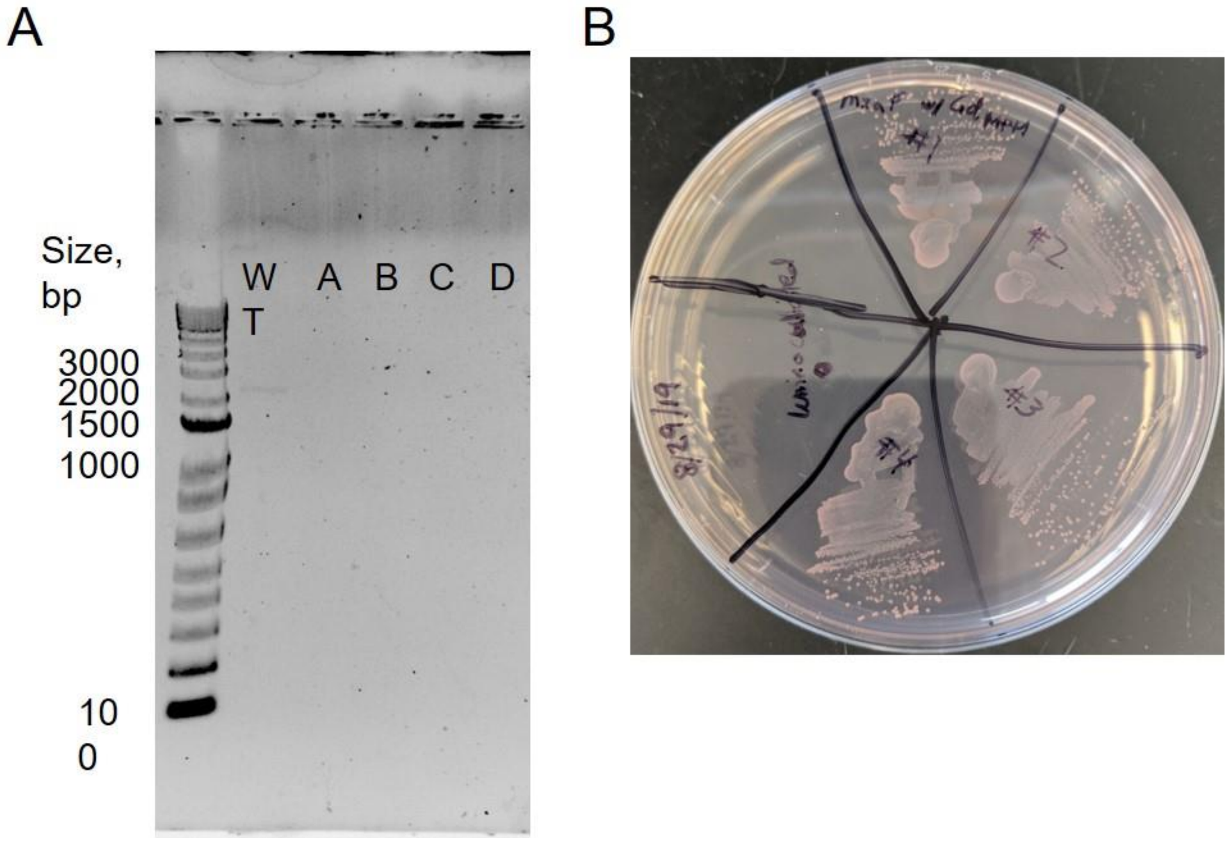


**FIG S1** PCR and phenotypic verification of the *mxaF* deletion in the *evo*-HLn strain recovered from methanol and $Gd^{3+}$ grown culture. *A*, Gel electrophoresis showing amplification of *mxaF* from wild type *M. extorquens* AM1 (WT) and no amplification from DNA of four independently grown *evo*-HLn cultures (A-D). PCR primers were designed to amplify ~500 bp upstream and downstream of *mxaF*. *B*, growth of four independently grown *evo*-HLn cultures (#1-4) on MP minimal medium with succinate and 50 µg/mL rifamycin after incubation at 30°C for 4 days. Medium from uninoculated growth medium incubated under the same conditions was spotted as a negative control.

**FIG S2**

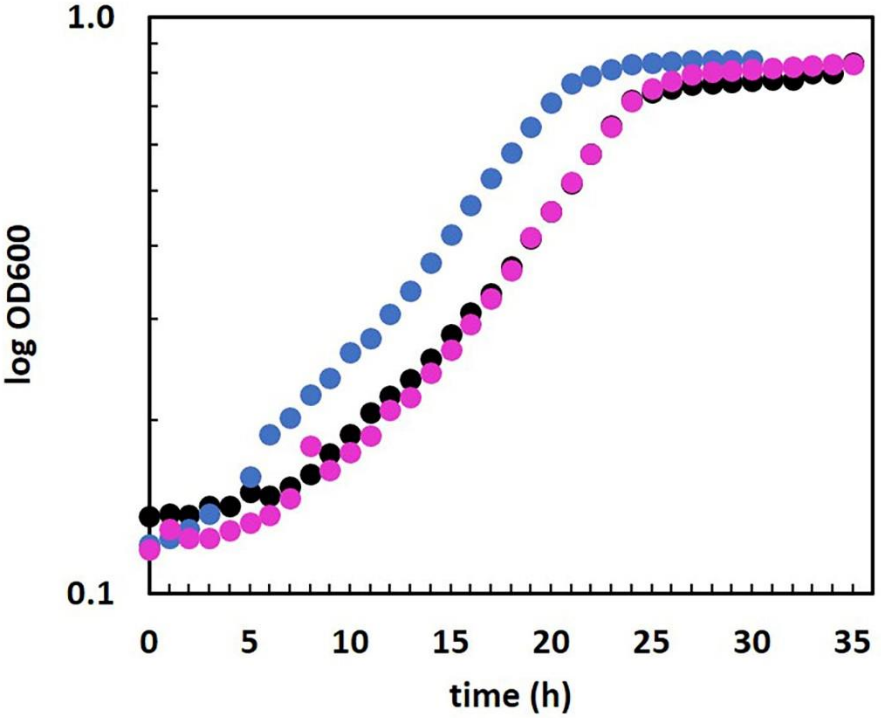


**FIG S2** Methanol growth of wild type *M. extorquens* AM1 with either no $Ln^{3+}$ (black), 2 μM $Gd^{3+}$ (pink) or 2 μM $La^{3+}$ (blue). Data points are the mean of 10 biological replicates from at least 3 independent experiments. Individual measurements for each data point are within 5% of one another.
